# PEPlife2: A Updated Repository of the Half-life of Peptides

**DOI:** 10.1101/2025.05.13.653654

**Authors:** Urooj Alam, Kunal Chaudhary, Nishant Kumar, Ritu Tomer, Sumeet Patiyal, Gajendra P. S. Raghava

## Abstract

This manuscript presents an updated version of PEPlife, a manually curated database that provides comprehensive information on peptide half-life. The updated version, PEPlife2, includes 4,412 entries, compared to 2,229 entries in the previous version. Each entry contains detailed information that include experimental methods used to determine half-life, chemical modifications, biological activity, routes of administration, and other relevant data. The database encompasses 402 proteins and 1,781 peptides, derived from 105 unique assays. In addition to natural peptide sequences, PEPlife2 includes cyclic peptides and chemically modified peptides, such as those with N- and C-terminal modifications. To provide structural insights, peptide and protein structures are sourced from the Protein Data Bank (PDB) or predicted using PEPstrMOD. To support the scientific community, we have developed a user-friendly interface and integrated advanced analytical tools, including BLAST, Smith-Waterman, GGSEARCH, CLUSTALW, and MUSTANG. The updated database is accessible at: https://webs.iiitd.edu.in/raghava/peplife2/.

**Author’s Biography:**

1. Urooj Alam is currently pursuing a Master’s degree in Computational Biology at the Department of Computational Biology, Indraprastha Institute of Information Technology, New Delhi, India.
2. Kunal Chaudhary is working as Ph.D. in Computational biology from Department of Computational Biology, Indraprastha Institute of Information Technology, New Delhi, India
3. Nishant Kumar is currently working as Ph.D. in Computational biology from Department of Computational Biology, Indraprastha Institute of Information Technology, New Delhi, India
4. Ritu Tomer is currently working as Ph.D. in Computational biology from Department of Computational Biology, Indraprastha Institute of Information Technology, New Delhi, India
5. Sumeet Patiyal is currently working as a postdoctoral visiting fellow Cancer Data Science Laboratory, National Cancer Institute, National Institutes of Health, Bethesda, Maryland, USA.
6. Gajendra P. S. Raghava is currently working as Professor and Head of Department of Computational Biology, Indraprastha Institute of Information Technology, New Delhi, India.

## Introduction

Peptide based therapeutics is the immensely growing field, whose market growth evaluated to be around USD 45.67 BN in 2023 and is projected to witness massive growth at a CAGR of around 5.63 % from 2024 to 2032 (USD 80.4 BN) [1]. The increasing market of peptide therapeutics and number of FDA approved peptide-based drugs shows the importance of peptide studies. Till now, 85 peptide based drugs have been approved by the FDA [2]. Around 150 peptides are currently in active clinical trials and with 260 peptides currently being tested in humans and 400 peptides in preclinical stages of development [3]. Despite these increasing demand of therapeutic peptides, their effectiveness is often limited by their short half-life due to their susceptibility to enzymatic degradation that reduces their bioavailability. Various experimental and in-silico applications have been utilize to determine half-life of peptides[4–10].

Most peptides exhibit a short *in vivo* half-life (t1/2 of 2–30 minutes), primarily due to enzymatic breakdown by proteases and rapid elimination through renal filtration, as molecules smaller than 30 kDa are swiftly excreted. Consequently, prolonging the *in vivo* half-life of peptides is crucial to maximize their therapeutic potential while reducing the need for high doses and frequent administration [11]. Attaching proteins to polyethylene glycol (PEG) (approximately 20–40 kDa) has been shown to effectively prolong their *in vivo* half-life. Alternatively, genetic fusion with antibody Fc domains or human serum albumin (HSA; 67 kDa) offers a promising strategy to replace PEGylation [12]. Additional techniques to enhance peptide stability and half-life include cyclizing peptides, substituting L-amino acids with D-amino acids, and creating hybrid peptides [11]. Physical modifications, such as coating amphiphilic peptides (AAPs) onto gold nanorods (GNRs), have also been utilized. This method extended the blood circulation half-life of GNRs in mice to 37.8 hours, significantly longer than the 26.6 hours observed with PEG-coated GNRs [13]. Thus, it is imperative to focus on this important topic, a large number of experimental and in-silico studies have been dedicated to improve and optimize the half-life of peptides [14–18].

In 2016, we developed the PEPlife database, which contains 2,229 entries, including 1,193 unique peptides and 37 unique proteins. This database was created to compile information on peptides whose half-life has been experimentally determined [16]. Over the past decade, PEPlife has been widely used and cited in scientific community. Additionally, datasets derived from PEPlife have been utilized to develop methods for predicting half-life of peptides [14,17]. Despite its extensive use by the scientific community, the database has not been updated since 2016. Therefore, there is a strong need to update it for the benefit of researchers working in the field of peptide and protein therapeutics. This manuscript introduces PEPlife2, an updated version of the original PEPlife database.

## Material and Methodology

### Data Curation

The PEPlife database, originally created in 2015, has been updated to gather new information from 2016 onwards to include new data that were not part of the previous version. We have collected the full text of granted patents from The Lens’s patents, and PubMed to search research publications. To gather the information from PubMed (https://pubmed.ncbi.nlm.nih.gov/), the query “((half-life[Title/Abstract]) AND (peptide[Title/Abstract]) NOT Review” was used to find pertinent publications on peptide half-life from the period between May 2016 and November 2024. The data was manually collected from research articles and patents. Only peptides with half-life values that were tested in experiments were included in the database. We have included new peptides (natural or synthesized/modified), tests, species, media, physical modification, and various half-life in this update. Initially, we obtained 1243 articles as on June 2024. During the initial screening, we exclude the articles and reviews that lack relevant information. We scrutinized around 950 articles to mine the relevant fields. Finally, we systematically curated the dataset from 449 articles. Additionally, we searched for the patents available in “The Lens” (https://www.lens.org/). It resulted in 18 patents as on June 2024. After that, we manually screened the 18 patents to retrieve the relevant information for data curation.

### Database Architecture and Web Interface

The standard platform “Linux-Apache-MySQL-PHP (LAMP)” was used to develop the PEPlife2 database. To set up the backend of the database, we used Apache (version 2.4.7) and MySQL (version 5.7.31) as the HTTP server. The front end of the webserver working with smartphones, tablets and desktops is developed using HTML (version 5), CSS (version 3), PHP (version 7.3.21) and JavaScript (version 1.8). A standard interface was created using the PHP programming language. The working architecture of PEPlife2 is explained in Figure 1.

**Figure 1.**
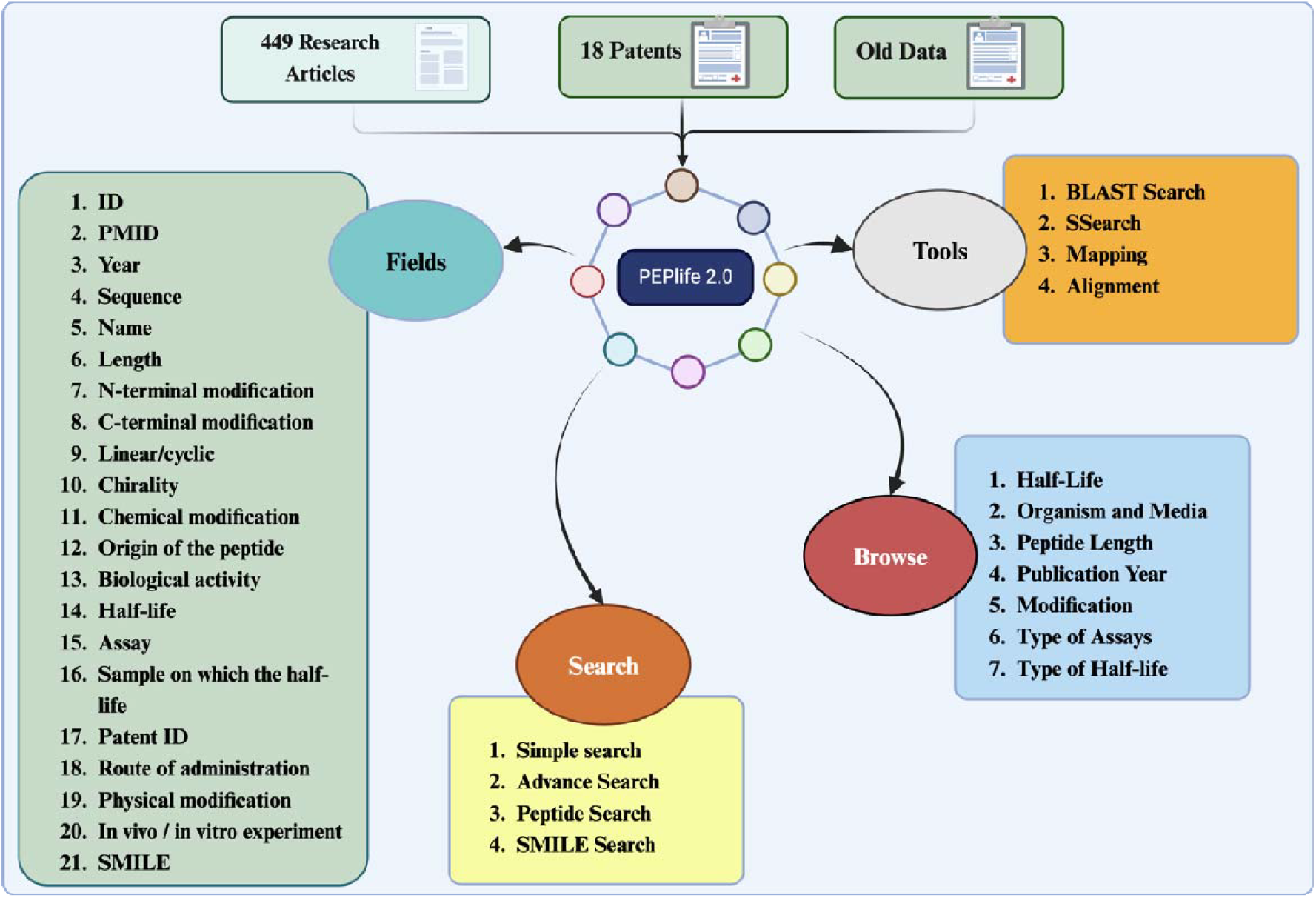
Architecture of peplife2

### Data Content

PEPlife2 contains information about peptide and protein half-life along with other fields which include: (1) PMID, (2) sequence, (3) name of peptide, (4) length, (5) N-terminal modification, (6) C-terminal modification (7) linear or cyclic structure (8) chirality, (9) chemical modification, (10) origin of the peptide, (11) biological activity of the peptide, (12) half-life, (13) assay types, (14) sample on which the half-life was tested and (15) Patent ID (16) route of administration (17) Physical modification (18) SMILES.

We have used these peptide/protein sequences for secondary information like tertiary structure and SMILES. In PEPlife2, peptide and protein structures were represented by 2D images. These structures were determined by searching the Protein Data Bank (PDB), assigning a structure if an exact match was identified. The PEPstrMOD webserver [19] was used to predict structures for peptides longer than five amino acids, in case there PDB structures were not available. Complex sequences with modifications like hydrocinnamate or chondroitin sulfate were excluded from structure prediction due to a lack of force field parameters. For peptides exceeding 40 residues, structure prediction was conducted using I-TASSER [20]. For peptides shorter than five residues, a linear conformation was adopted with dihedral angles (□and ψ) set to 180°, followed by energy minimization and molecular dynamics (MD) simulations for refinement. Secondary structures of all peptides were derived from their tertiary structures using DSSP (Define Secondary Structure of Proteins) software [21], which classifies each residue into one of eight states: Beta-bridge (B), Loop (C), Extended strand (E), 3/10 helix (G), Alpha-helix (H), Pi-helix (I), Bend (S), and Turn (T).

### Browsing Tool

The PEPlife2 browsing module make it easy to retrieve data through the class-wise browsing feature, in which all the peptide entries are classified into different categories, include “Half-life”, “Organism and Media”, “Peptide Length”, “Publication Year”, “Type of Modification”, “Type of Assay”.

### Analysis Tool

Various bioinformatics tools have been integrated to analyze query sequences against PEPlife2 entries. The Basic Local Alignment Search Tool (BLAST) and Smith-Waterman used to enable similarity-based search[22–24]. The “Peptide Mapping” was included for peptide similarity analysis and peptide mapping. The “Sequence Alignment” page aligns query sequence with PEPlife2 entries. Sequence Alignment was done using CLUSTAL [http://www.clustal.org/omega/] tool. Another page provides “Structural Alignment” by using a query PDB file and aligning it against PEPlife2 PDB structures using the MUSTANG tool [25].

## Result and Discussion

The PEPlife2 (https://webs.iiitd.edu.in/raghava/peplife2/) is an updated version of the PEPlife database, containing a total of 2,183 entries alongwith 74 miscellaneous entries, comprising 2,046 entries collected from 449 research articles and 211 entries extracted from 18 patents. The database includes peptides of various lengths, ranging from single-residue peptides to those with more than 54 residues. The group of peptides with length ranges from 1 to 10 residues represents the largest category, followed by peptides in the 11 to 21 residue range, and then those in the 33 to 43 residue range, while the 44 to 54 residue group has the smallest number of entries. Structurally, most peptides in the database are linear, accounting for 3,728 entries, compared to 655 entries representing cyclic peptides. Additionally, terminal modifications are common, with 2,104 peptides having N-terminal modifications and 1,481 peptides featuring C-terminal modifications. Various assays were utilized to measure the half-life of the peptides, with mass spectrometry being the most commonly employed method, followed by HPLC and ELISA. A detailed distribution of peptides based on their lengths, D/L-configuration (or mixed configuration), structural conformations, and the assays used to evaluate their half-life is presented in Figure 2.

**Figure 2.**
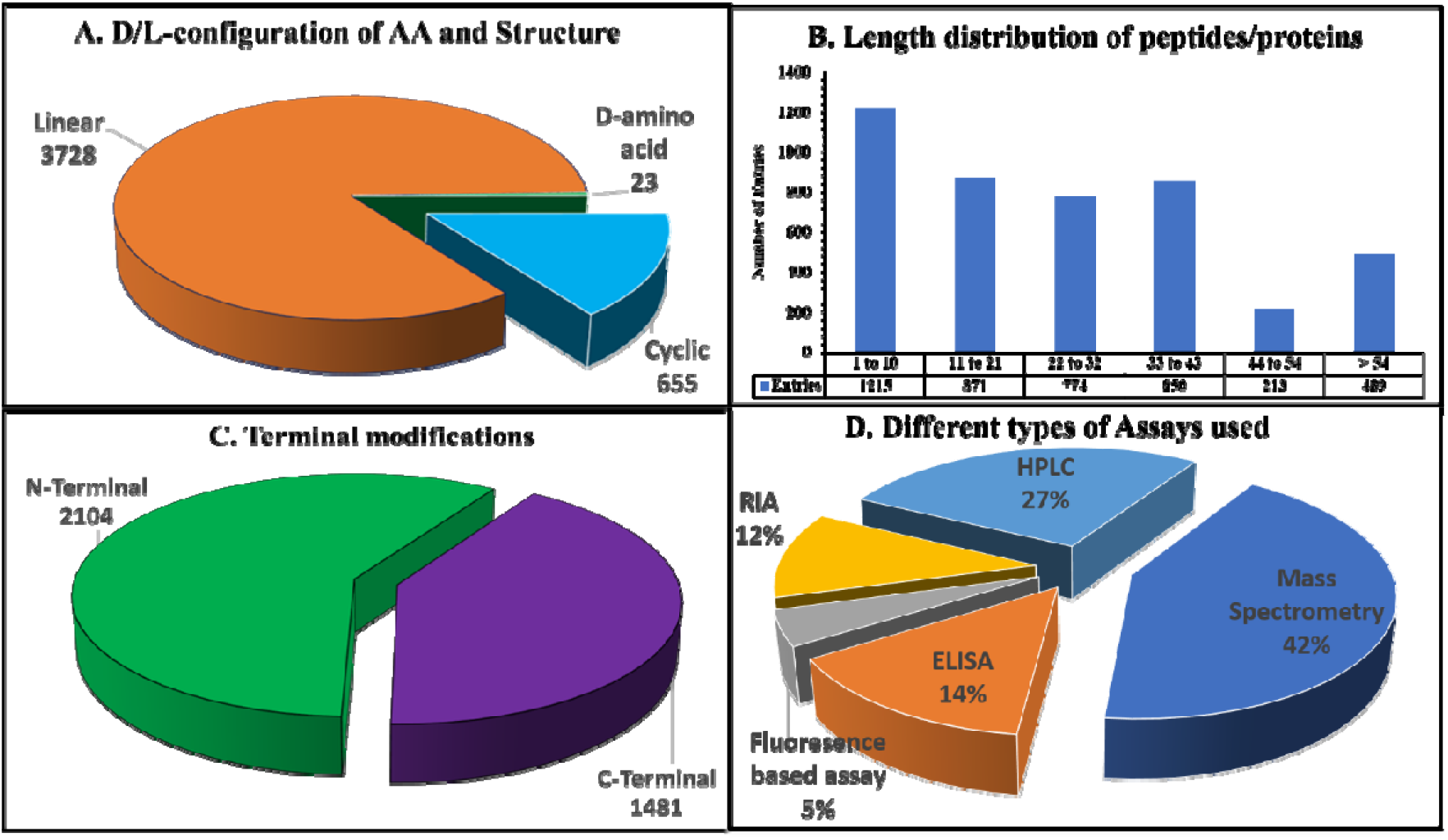
Distribution of peptides/proteins based on (A) Length (B) D/L-configuration of amino acids and Structure, (C)Terminal modifications, (D) Different types of Assays use

Also, a detailed distribution of unique peptides and protein length distribution is given below in Figure 3.

**Figure 3.**
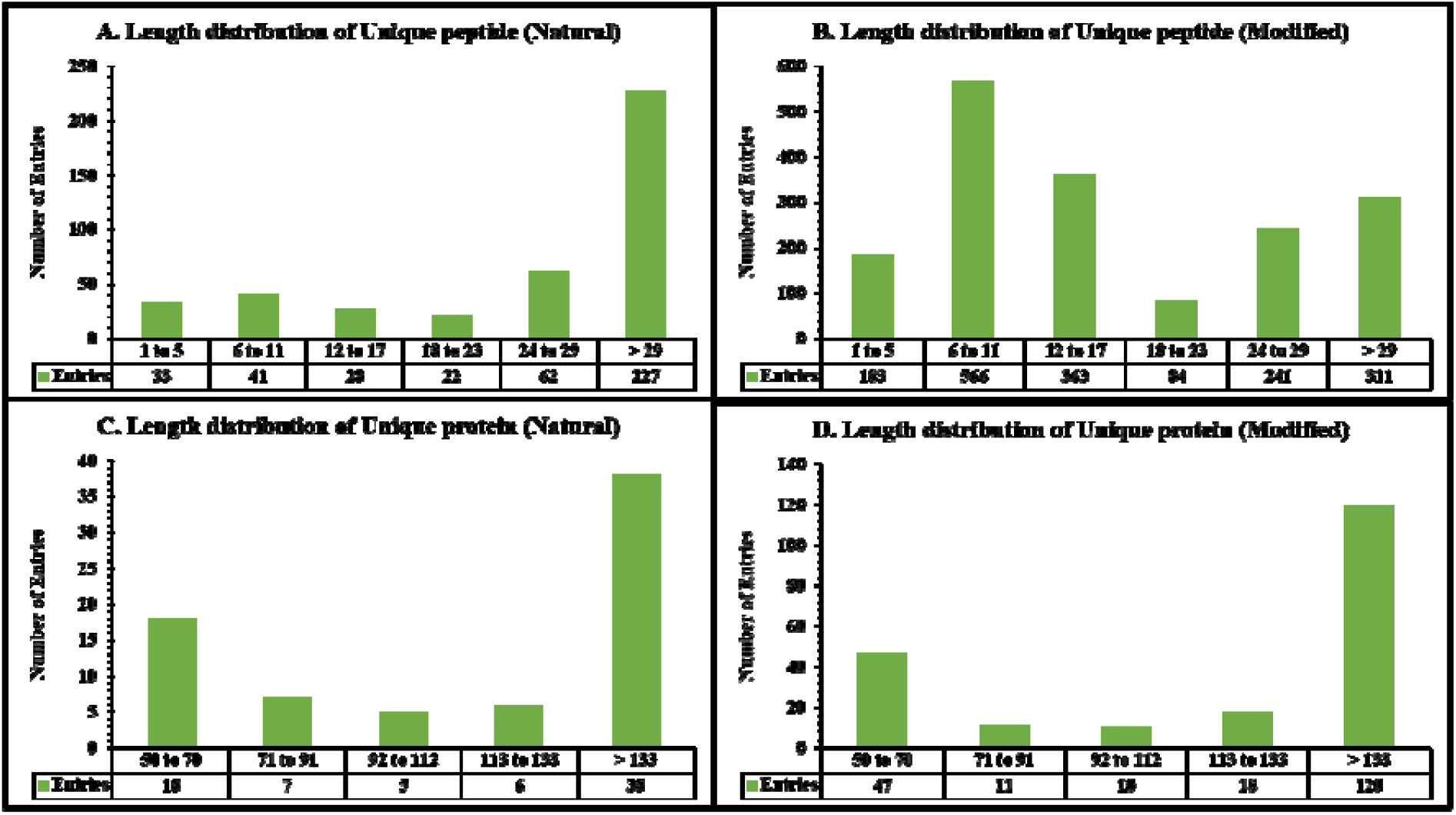
Length distribution of Unique peptides and proteins

Our database includes various chemical modifications aimed at enhancing the stability of peptides or proteins, such as PEGylation, Fc-conjugation, amino acid replacements (e.g., Lys substitutions followed by fatty acid conjugation), D-amino acid substitutions, non-natural amino acid substitutions (e.g., Aib, Sar, pGlu), cyclization, amidation, acetylation, and others. The effects of these chemical modifications are detailed in Figure 4. For example, physical modification of E1 NPs (polymeric nanoparticles coated with GC containing the HIV-1 fusion inhibitor peptide E1) extended its half-life from 8 hours to 24 hours compared to its natural form, E1. Similarly, terminal modifications like N-terminal acetylation and C-terminal amidation significantly improved the half-life of peptide Ang(1-7), increasing it from approximately 9 minutes to 135 minutes when compared to its natural, unmodified form. Incorporating D-amino acids into VH445 enhanced its half-life from 1.16 hours to 3.03 hours compared to the unmodified VH434. Furthermore, unnatural amino acids, such as Aib and Nle, also contributed to longer half-life, with Nle extending the half-life up to 29 hours. Additionally, we have incorporated the MAP (Modification and Annotation in Proteins) [26] format that introduces tags within the sequence for residue-level modification such as chemical modifications, non-standard amino acids, binding sites, mutations.

**Figure 4.**
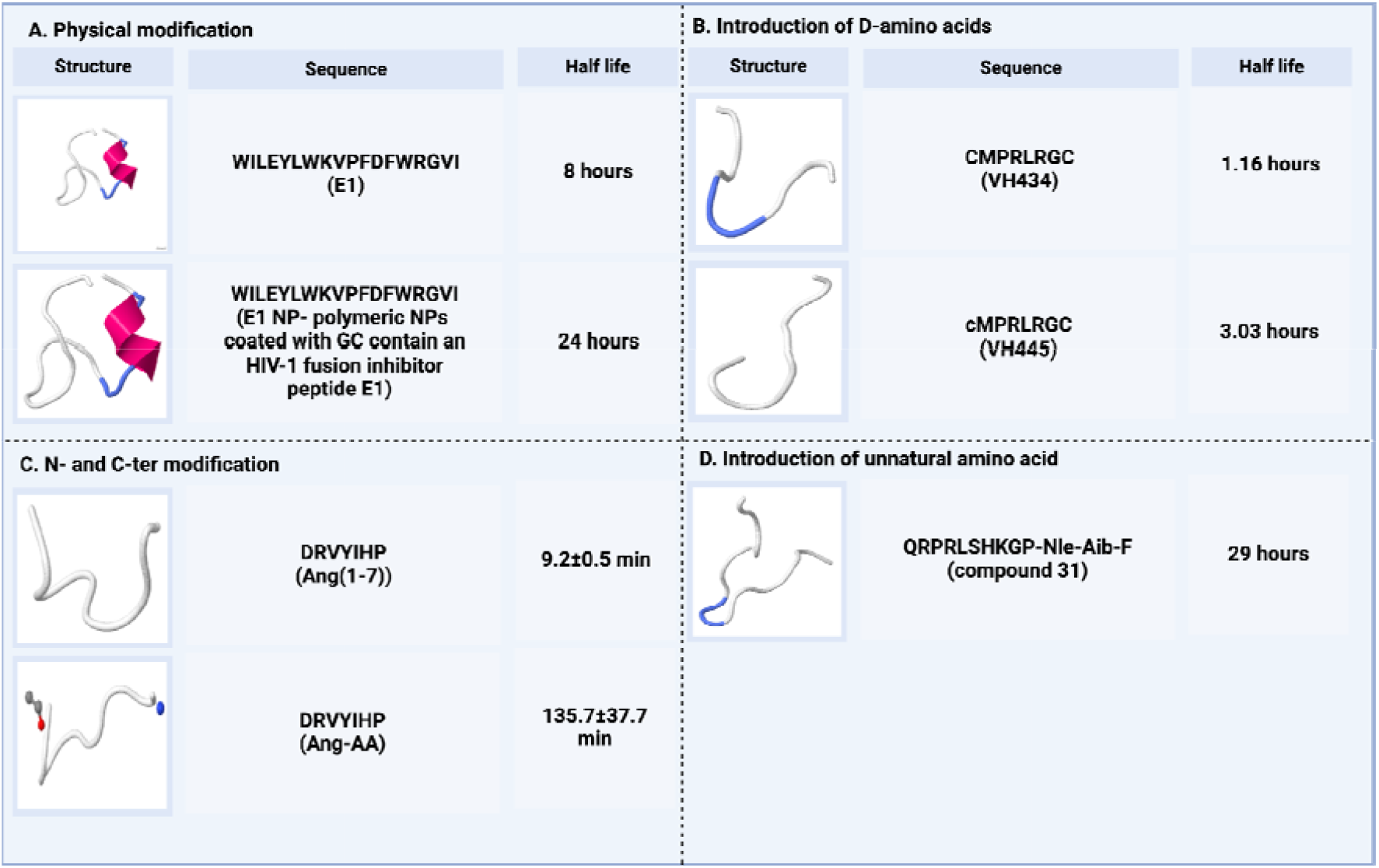
Effect of various modifications on peptide or protein half life

### Comparison with previous PEPlife version

The original PEPlife database (published in 2016, covering data up to 2015) comprised 2,229 entries, representing 1,193 unique peptides. Each entry provided detailed information, including the peptide’s name, sequence, half-life, modifications, the experimental assay used to determine half-life, as well as its biological nature and activity.

In the updated version, entries from 2016 to 2024 have been added, sourced from 449 research articles and 18 patents, significantly expanding the database. This new version introduces several enhancements that were absent in the original, such as physical modifications (e.g., hydrogels), which are now commonly employed to increase peptide half-life. Here below is the comparison table between PEPlife and PEPlife2 statistics for different fields in Figure 5.

**Figure 5.**
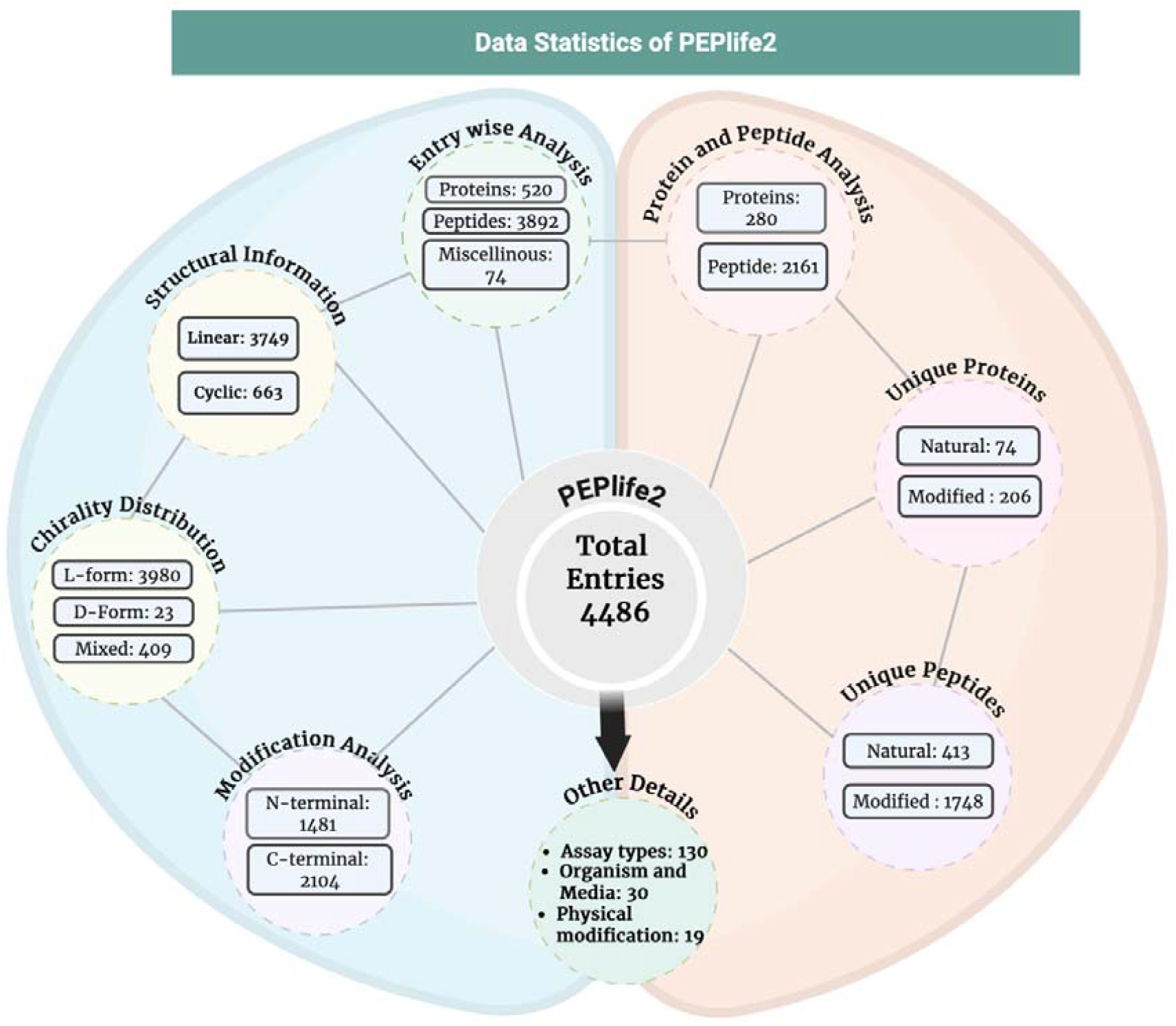
The complete statistics of PEPlife2 data

The key improvements in functionality include the elimination of the requirement to install Jalview (previously needed for sequence and structure alignment). Users can now view results directly on specific pages, streamlining the process. Additionally, the updated version includes data for various types of half-life (such as distribution half-life and activity half-life) and physical modification for more effective categorization and exploration of these types. These updates significantly enhance the usability of the database, making it a more comprehensive resource for understanding peptide stability and modifications compared to the original version.

## Conclusion

We have created an updated version, PEPlife2, which includes 2,183 new entries along with 74 miscellaneous ones, in addition to the 2,229 existing entries from PEPlife. This update expands the dataset to a total of 4,412 entries. The PEPlife2 database is available at PEPlife-2 (https://webs.iiitd.edu.in/raghava/peplife2/). This provides better coverage of all peptide and protein half-life data. Recently discovered half-life peptides and proteins from the most recent research have been added to the database, updating the data. Either getting the data directly from the website or accessing it programmatically makes it simple to access. We think the scientific community will benefit from the updated edition.

### Limitations & update of peplife2

We have made a concerted effort to provide comprehensive information on the half-life of proteins and peptides; however, certain peptides could not be included due to their absence in patents or other publications. Additionally, the unavailability of force field libraries for complex chemical modifications resulted in some structures being excluded. Despite meticulous data analysis, absolute precision cannot be guaranteed, as the possibility of human error remains. With the release of PEPlife2 (https://webs.iiitd.edu.in/raghava/peplife2/), we aim to enhance both the quality and scope of the database to assist the scientific community.

## Availability

The database PEPlife2 is available free of cost at “https://webs.iiitd.edu.in/raghava/peplife2/”, and the data can be downloaded from the downloads tab provided on the website or directly from the API by selecting the query field at https://webs.iiitd.edu.in/raghava/peplife2/api/rest.html.

## Authors’ contributions

UA, GPSR, and NK manually collected the data. UA, KC, and RT manually curated and analysed the data. NK, SP, and UA developed the backend and frontend of the webserver. UA, NK, and GPSR prepared the manuscript. NK, UA, KC, RT, SP, and GPSR. reviewed the manuscript. GPSR conceived and coordinated the project. All authors read and approved the final manuscript.

## Acknowledgments

Authors are thankful to the University Grants Commission (UGC), Department of Science and Technology (DST-INSPIRE), for fellowships and financial support, Indraprastha Institute of Information Technology (IIITD), for fellowships and financial support, and the Department of Computational Biology, IIITD, New Delhi, for infrastructure and facilities. We would like to acknowledge that Figures were created using BioRender.com.

